# ALGAEFUN with MARACAS, microALGAE FUNctional enrichment tool for MicroAlgae RnA-seq and Chip-seq AnalysiS

**DOI:** 10.1101/2021.08.14.456338

**Authors:** Ana B. Romero-Losada, Christina Arvanitidou, Pedro de los Reyes, Mercedes García-González, Francisco J. Romero-Campero

## Abstract

**Background:** Microalgae are emerging as promising sustainable sources for biofuels, biostimulants in agriculture, soil bioremediation, feed and human nutrients. Nonetheless, the molecular mechanisms underpinning microalgae physiology and the biosynthesis of compounds of biotechnological interest are largely uncharacterized. This hinders the development of microalgae full potential as cell-factories. The recent application of omics technologies into microalgae research aims at unraveling these systems. Nevertheless, the lack of specific tools for analysing omics raw data generated from microalgae to provide biological meaningful information are hampering the impact of these technologies. The purpose of ALGAEFUN with MARACAS consists in providing researchers in microalgae with an enabling tool that will allow them to exploit transcriptomic and cistromic high-throughput sequencing data.

**Results:** ALGAEFUN with MARACAS consists of two different tools. First, MARACAS (MicroAlgae RnA-seq and Chip-seq AnalysiS) implements a fully automatic computational pipeline receiving as input RNA-seq (RNA sequencing) or ChIP-seq (chromatin immunoprecipitation sequencing) raw data from microalgae studies. MARACAS generates sets of differentially expressed genes or lists of genomic loci for RNA-seq and ChIP-seq analysis respectively. Second, ALGAEFUN (microALGAE FUNctional enrichment tool) is a web-based application where gene sets generated from RNA-seq analysis as well as lists of genomic loci from ChIP-seq analysis can be used as input. On the one hand, it can be used to perform Gene Ontology and biological pathways enrichment analysis over gene sets. On the other hand, using the results of ChIP-seq data analysis, it identifies a set of potential target genes and analyses the distribution of the loci over gene features. Graphical representation of the results as well as tables with gene annotations are generated and can be downloaded for further analysis.

**Conclusions:** ALGAEFUN with MARACAS provides an integrated environment for the microalgae research community that facilitates the process of obtaining relevant biological information from raw RNA-seq and ChIP-seq data. These applications are designed to assist researchers in the interpretation of gene lists and genomic loci based on functional enrichment analysis. ALGAEFUN with MARACAS is publicly available on https://greennetwork.us.es/AlgaeFUN/.

## Background

Microalgae are a very diverse non-monophyletic group of photosynthetic microorganisms of special interest due to their physiological plasticity and wide range of biotechnological applications. Microalgae can be found in a wide variety of different habitats, from freshwater to oceans growing under a broad range of temperature, salinity, pH and light intensity values. More than 5000 species have been identified in the oceans accounting for the production of 50% of the oxygen necessary to sustain life on Earth. Microalgae also play a central ecological role as primary producers of biomass establishing the base of aquatic food chains [1]. In the last decades, microalgae have also been of great interest for the scientific community due to the large and yet increasing number of biotechnological applications they present. Specifically, they have been described as a high yield source of carbon compounds and good candidates for the mitigation of CO2 emissions. In microalgae, fixation of CO2 is coupled with growth and biosynthesis of compounds of biotechnological interests such as polysaccharides, lipids, vitamins and antioxidants. Their industrial cultivation for the production of biofuels as well as bioproducts used as biostimulants in agriculture, health supplements, pharmaceuticals and cosmetics is also thoroughly explored nowadays [2,3]. Finally, microalgae are successfully applied in wastewater treatment coupled with fixation of atmospheric CO2 [4].

Due to these promising features, physiological characterization of multiple microalgae species under different cultivation regimes have been thoroughly carried out [5,6]. Nevertheless, the molecular mechanisms underpinning the physiology of microalgae are yet poorly understood. In order to facilitate the progress in the characterization of the molecular systems regulating microalgae physiology, high throughput sequencing technologies have been recently applied to obtain the genome of a wide range of microalgae [7–19]. This has promoted the use of different omics, particularly transcriptomics based on RNA-seq data [20–22] and cistromics based on ChIP-seq data [23,24], to initiate molecular systems biology studies in microalgae. Nonetheless, the impact of this type of studies on microalgae are hampered by the lack of freely available and easy to use online tools to analyze, extract relevant information and integrate omics data. Typically, RNA-seq and ChIP-seq analysis received as input high-throughput sequencing raw data and produce as output, respectively, sets of differentially expressed genes and genomic loci significantly bound by the protein of interest. Processing of the massive amount of high-throughput sequencing data and analysis of the resulting sets of genes and genomic loci obtained from molecular systems biology studies requires computational power, time, effort and expertise that research groups on microalgae may lack. In addition, researchers must explore different data bases separately, which makes the integration of the results and the generation of biological meaningful information more difficult.

Therefore, it is imperative the development of frameworks integrating microalgae genome sequences and annotations with tools for high-throughput sequencing data analysis and functional enrichment of gene and genomic loci sets. In order to cover these microalgae research community needs and promote studies in molecular systems biology we have developed the web portal ALGAEFUN with MARACAS using the R package Shiny [25] and other Bioconductor packages. Our web portal consists of two different tools. MARACAS (MicroAlgae RnA-seq and Chip-seq AnalysiS) implements an automatic computational workflow that receives as input RNA-seq or ChIP-seq raw sequencing data from microalgae studies and produces, respectively, sets of differentially expressed genes or a list of genomic loci. These results can be further analyzed using our second tool ALGAEFUN (microAlgae FUNctional enrichment tool). On the one hand, when receiving the results from an RNA-seq analysis, sets of genes are functionally annotated by performing GO (Gene Ontology) [26] and KEGG (Kyoto Encyclopedia of Genes and Genomes) pathways [27] enrichment analysis. On the other hand, when genomic loci from a ChIP-seq analysis are inputted, a set of potential target genes is generated together with the analysis of the distribution of the loci over gene features as well as metagene plots representing the average mapping signal. This set of potential target genes can be further studied using the features for functional enrichment analysis in ALGAEFUN as described above. ALGAEFUN with MARACAS supports a wide range of 14 different microalgae species including *Chlamydomonas reinhardtii*, *Ostreococcus tauri*, *Nannochloropsis gaditana* and *Phaeodactylum tricornutum* among others. Additionally, our tools can be easily extended to include new microalgae as their genome sequence and annotation are made available. The code for ALGAEFUN with MARACAS is publicly available at their respective GitHub repositories from the following links: https://github.com/fran-romero-campero/ALGAEFUN and https://github.com/fran-romero-campero/MARACAS.

### Implementation

ALGAEFUN with MARACAS supports the analysis of the following microalgae covering an ample spectrum of their phylogeny: *Chlamydomonas reinhardtii* [7], *Volvox carteri* [8], *Chromochloris zofingiensis* [9], *Dunaliella salina* [10], *Haematococcus lacustris* [11] (Chlorophyceae), *Coccomyxa subellipsoidea* [12] (Trebouxiophyceae), *Ostreococcus tauri* [13], *Bathycoccus prasinos* [14], *Micromonas pusilla CCMP1545* [15] (Mamiellophyceae), *Phaeodactylum tricornutum* [16], *Nannochloropsis gaditana* [17] (Stramenopiles), *Klebsormidium nitens* [18], *Mesotaenium endlicherianum* and *Spirogloea muscicola* [19] (Charophyceae), Fig. 1.

**Figure 1.**
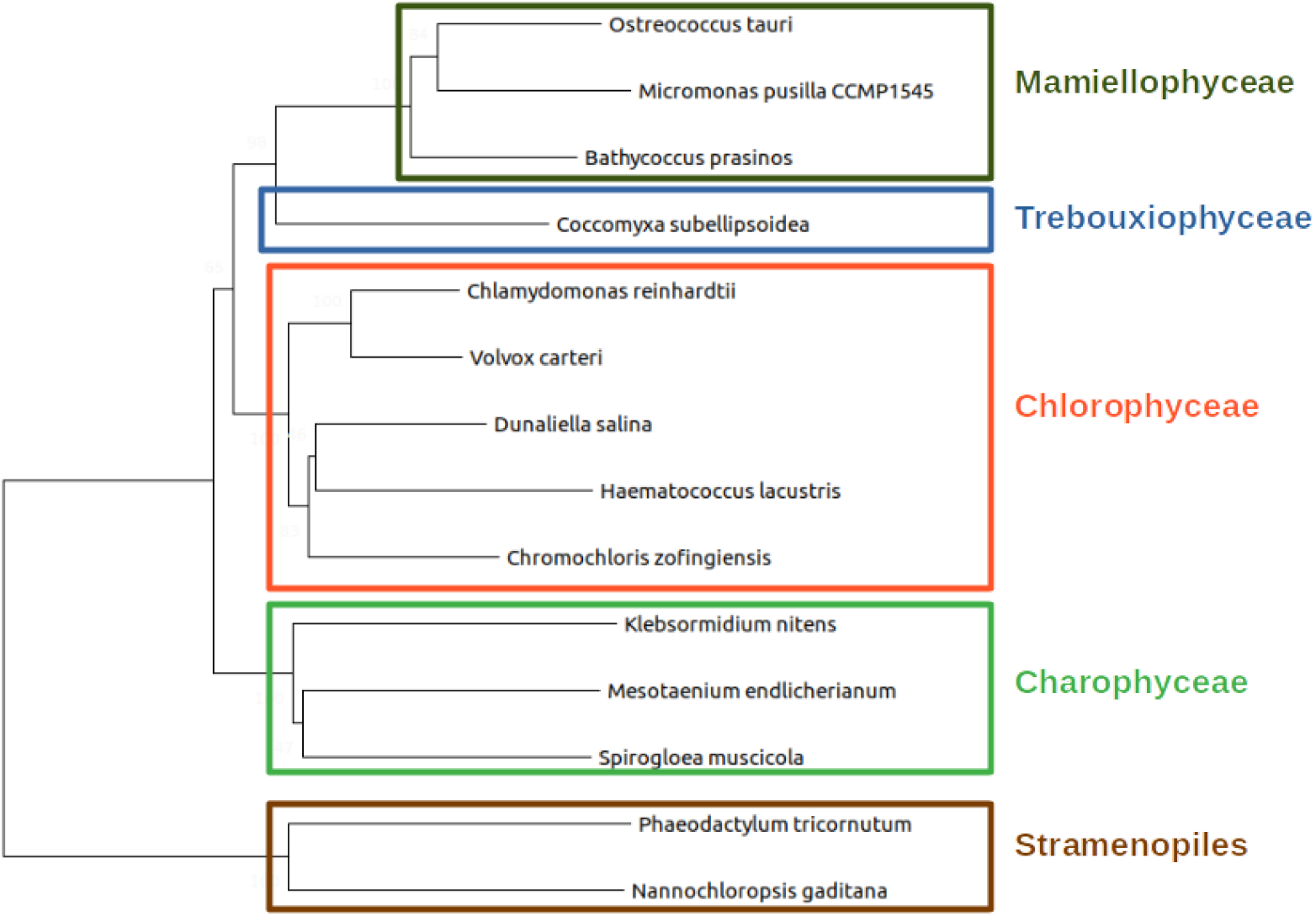
Phylogenetic relationship between the different microalgae species supported in ALGAEFUN with MARACAS. The Mamiellophyceae *Ostreococcus tauri*, *Micromonas pusilla CCMP1545* and *Bathycoccus prasinos* (olive green rectangle); the Trebouxiophyceae *Coccomyxa subellipsoidea* (blue rectangle); the Chlorophyceae *Chlamydomonas reinhardtii*, *Volvox carteri*, *Dunaliella salina, Haematococcus lacustris* and *Chromochloris zofingiensis* (red rectangle); the Charophyceae *Klebsormidium nitens*, *Mesotaenium endlicherianum* and *Spirogloea muscicola* (green rectangle); and the Stramenopiles *Phaeodactylum tricornutum*, *Nannochloropsis gaditana* (brown rectangle).

One of the limiting factors for the development of molecular systems biology studies in microalgae is the fragmentation of the available genomic sequences and functional annotations into different databases. To overcome this issue and generate easily accessible resources, genome sequences, functional annotation and genomic feature annotation files for the previous microalgae species were systematically collected from different freely available databases: Ensembl Protists [28], PhycoCosm [29], Phytozome [30], Genomes - NCBI Datasets [31] and a Figshare repository [32], see Table 1.

**Table 1.**
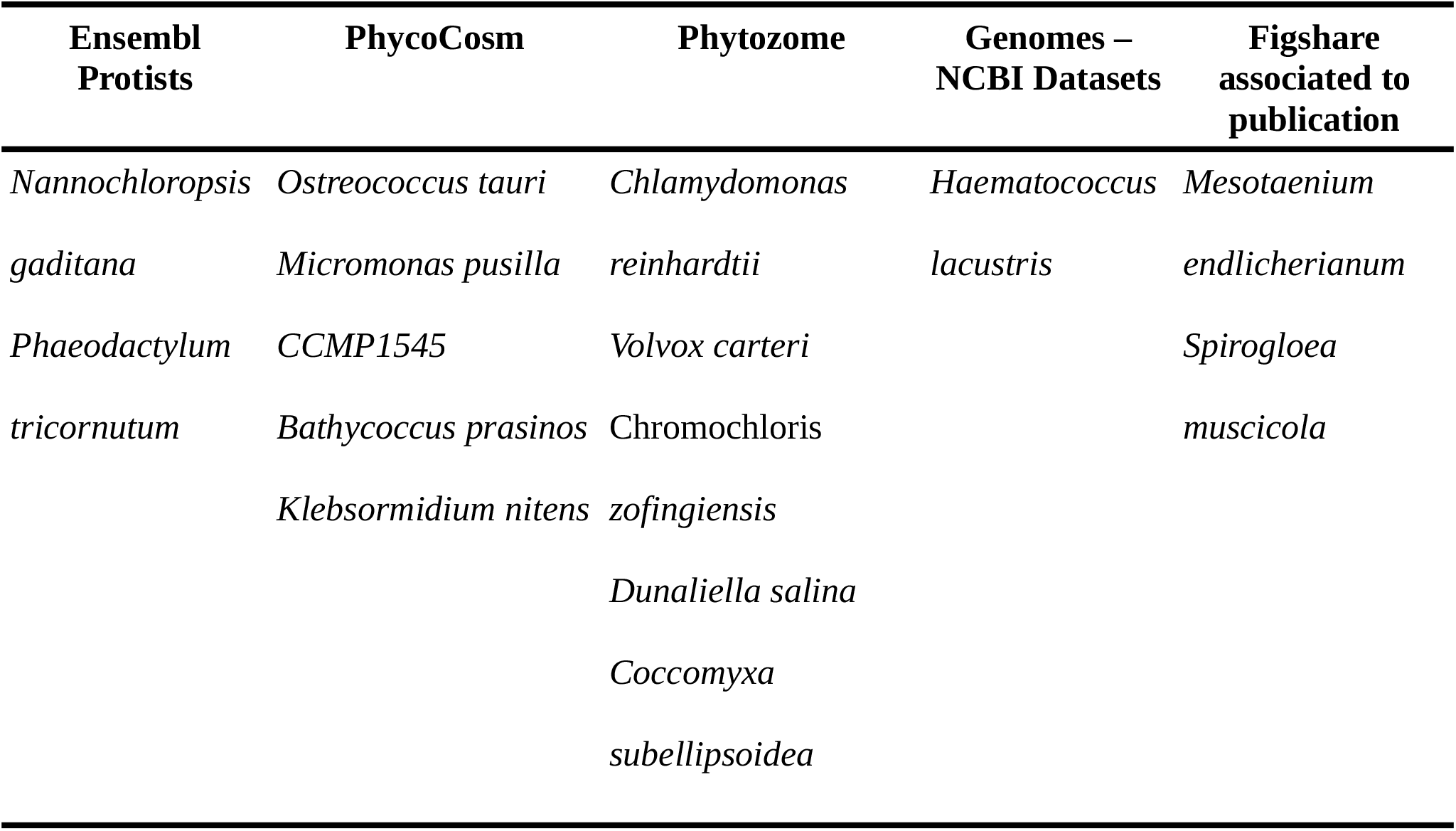
Resources used to collect genome sequences, functional and gene feature annotations for each supported microalga.

The MARACAS pipeline is based on the short read mappers HISAT2 (Hierarchical Indexing for Spliced Alignment of Transcripts 2) [33] and Bowtie2 [34]. Pre-computed genome indexes for each microalga are included in MARACAS to alleviate the high-cost computational tasks consisting of read mapping to reference genomes. Gene features annotation files in GTF (gene transfer format) for each microalga were also collected. Transcript assembly and estimation of gene expression is performed using StringTie [33]. Identification of differentially expressed genes and graphical representation of the results are carried out using the R Bioconductor packages Ballgown [33] and LIMMA (Linear Models for Microarray Analysis) [35]. For ChIP-seq data analysis, the peak caller MACS2 (Model-based Analysis of ChIP-seq 2) [36] is used. MARACAS can be executed in sequential mode or in distributed/parallel mode on a computational cluster managed by the job scheduling system SLURM (Simple Linux Utility for Resource Management). In the distributed/parallel mode, synchronization among the jobs processing different samples is implemented using a blackboard system. A common blackboard file is used where parallel jobs can read and write to ensure that all samples have been processed before proceeding to the next steps. This effectively implements synchronization points during the workflow in MARACAS.

For the implementation of ALGAEFUN, functional annotation files were also downloaded for each microalga from the previously mentioned databases, Table 1. Specifically, Gene Ontology (GO) and KEGG (Kyoto Encyclopedia of Genes and Genome) Orthology (KO) terms were collected. For microalgae species lacking these annotation systems HMMER (biological sequence analysis using profile hidden Markov models) [37] was used to identify protein domains according to the PFAM (Protein Family) nomenclature. PFAM terms were subsequently converted into GO terms using pfam2go. KO terms were associated to genes applying KAAS (KEGG Automatic Annotation Server) [38]. Additionally, whenever available other systematic functional annotation formats were also included such as Protein Analysis Through Evolutionary Relationships (PANTHER) terms [39], Enzyme Commission numbers (EC numbers) and Eukaryotic Orthologous Groups (KOG) terms [40]. In order to use all these functional annotation systems in ALGAEFUN two different types of R annotation packages were developed and were made freely available from our Github repository [41]. On the one hand, using the function makeOrgPackage from the Bioconductor R package AnnotationForge [42] we generated annotation packages for each microalga integrating the systematic sources of functional annotation discussed previously. These packages are instrumental when performing functional enrichment analysis over gene sets obtained, for instance, from a differential expression analysis based on RNA-seq data using MARACAS. On the other hand, applying the function makeTxDbFromGFF from the Bioconductor R package GenomicFeatures [43] we developed a package for each microalga storing gene features annotation available from the previously downloaded and processed GTF files. These packages are central to carry out analysis over genomic loci obtained, for example, using MARACAS and ChIP-seq data to study the genome wide occupation of specific transcription factors or histone modifications. These genomic and functional annotation packages will enable the microalgae research community to perform omics analysis independently from the tools available in ALGAEFUN with MARACAS. The interactive interface of ALGAEFUN was developed using the R package Shiny [25]. In turn, ALGAEFUN functionalities were implemented based on our annotation packages described above. The R Biocondutor packages clusterProfiler [44] and pathview [45] are used for functional enrichment analysis over gene sets. Whereas the annotation and visualization of genomic loci is performed in ALGAEFUN with the R Bioconductor packages ChIPseeker [46] and ChIPpeakAnno [47].

## Results and Discussion

Our web-based applications ALGAEFUN (MicroALGAE FUNctional annotation tool) with MARACAS (MicroAlgae RnA-seq and Chip-seq AnalysiS) seek to become enabling tools that would promote molecular systems biology studies in microalgae. A wide range of microalgae species relevant both in basic research and biotechnological applications are supported. Moreover, our tools are easily extendable to include new microalgae species as their genome sequences and functional annotations become available. Analysis combining the different functionalities implemented in our tools will allow researchers to extract relevant information in the form of functional annotation enrichment analysis over gene sets and genomic loci starting from raw high-throughput sequencing data. ALGAEFUN with MARACAS has been applied recently to determine the molecular systems underpinning astaxanthin accumulation in the microalgae of industrial interest *Haematococcus lacustris* [21].

Next, we use two case studies starting from RNA-seq and ChIP-seq raw sequencing data respectively to describe the user interface and discuss the intended uses and benefits of applying ALGAEFUN with MARACAS in microalgae research.

### Case Study 1. From RNA-seq raw sequencing data to biological processes and pathways

A detailed description of the steps to install and execute MARACAS is provided at the corresponding Github repository [48] and the video tutorials available on our webpage [49]. Using the automatic pipeline maracas-rna-seq, users can process raw high-throughput sequencing data in fastq format available either locally in their computers or from a data base such as Gene Expression Omnibus [50] or Sequence Read Archive [51]. This pipeline can be executed either in sequential mode or in distributed/parallel mode on a computational cluster managed by the job scheduling system SLURM. The parameter settings need to be provided in a single text file that constitutes the only input to this pipeline. Besides specifying the microalga of interest, the execution mode and fastq files locations or accession numbers, these parameters describe the experimental design, control and experimental samples, as well as the fold-change and significance level thresholds for the identification of differentially expressed genes. Reports in html and pdf format are generated containing information regarding sequence quality analysis, mapping process and differential gene expression.

In order to illustrate this pipeline, we re-analysed RNA-seq data studying the response of the mamiellophyceae microalgae *Ostreococcus tauri* to iron starvation [20]. The parameter file to reproduce this analysis is provided within the MARACAS distribution bundle. The reports produced by MARACAS described all samples as of high quality and notified no problem during read mapping to the reference genome with mapping rates greater than 94%. Scatter plots comparing gene expression between samples are also produced in the MARACAS report. In this case, high Pearson correlations greater than 98% were identified between replicates of the same condition. Accordingly, the automatically performed Principal Components Analysis in MARACAS identified two clearly separated clusters constituted by the control and iron starvation samples, Fig 2. A volcano plot is used in the report to represent the 45 repressed genes and 554 activated genes identified with a fold-change threshold of two and a q-value threshold of 0.01, Fig 2. These lists of genes can be then inputted into ALGAEFUN to determine significantly overrepresented biological processes or pathways affected during iron starvation in *Ostreococcus*.

**Figure 2.**
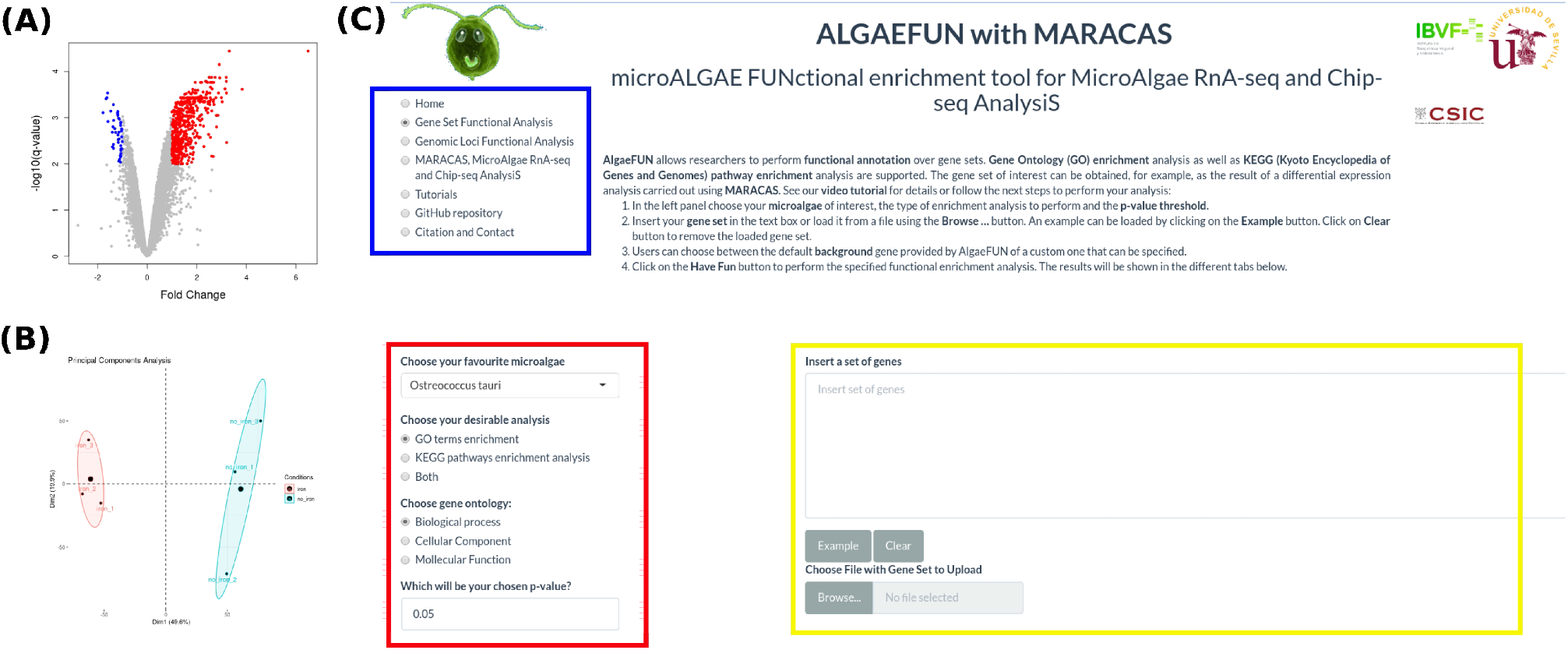
RNA-seq MARACAS results and Graphical Interface in AlgaeFUN to perform Gene Set Functional Enrichment Analysis. (A) Volcano plot automatically generated by MARACAS marking activated genes in red and repressed genes in blue. (B) Principal Components Analysis automatically generated by MARACAS clearly separating samples into two different clusters depending on the analysed conditions. (C) AlgaeFUN Graphical Interface for Functional Enrichment Analysis, navigation side panel for selecting the different functionalities (blue box); dropdown menu for microalgae selection and sidebar panel with parameter settings for GO and/or KEGG pathways enrichment analysis (red box); text box for gene set input and gene background selections (yellow box).

The graphical interface in ALGAEFUN allows users to explore the different functionalities from a navigation side panel, Fig. 2(C). For this case study we selected “Gene Set Functional Analysis”. Next, the microalga of interest needs to be chosen from a dropdown menu. Here, we chose *Ostreococcus tauri*. Users need to specify whether GO term and/or KEGG pathway enrichment analysis are to be performed at a selected significance level. The set of genes to analyse can be specified through a text box or by uploading the corresponding file. As background gene set, users can also choose between the entire microalga genome selected by default or provide a custom gene set. ALGAEFUN provides gene set examples for each microalga so that users can explore our tool and check the required gene id format. These examples can be accessed and inputted in the corresponding text box by clicking on the Example button. These examples were generated during the testing of MARACAS using previously published RNA-seq data sets and have in turn been used in the testing and validation of ALGAEFUN. In this case study we decided to carry out a GO term and KEGG pathway enrichment analysis over the set of 554 activated genes under iron starvation using the default background gene set for *Ostreococcus*.

The outputs of the functional enrichment analysis are presented in the graphical interface in different tabs. Downloadable tables are generated consisting of columns with GO/KEGG term identifiers, human readable descriptions, p-values, q-values, enrichment values and the list of genes associated in the input set with the corresponding GO/KEGG term. Gene names can be clicked to access their annotation from different data bases. Furthermore, ALGAEFUN also generates several graphs that represent the GO/KEGG term enrichment that can be visualized and downloaded from the different tabs such as acyclic graphs, barplots, dotplots, enrichment maps, gene-concept networks and KEGG pathway maps. Our results were in agreement with the published ones [20] identifying ribosome biogenesis and DNA metabolic process as key biological processes overrepresented in the set of activated genes.

### Case Study 2. From ChIP-seq raw sequencing data to marked genes

Similar to the previous case study, MARACAS contains an automatic pipeline that can be executed in sequential or parallel mode, maracas-chip-seq. This pipeline starts from raw high-throughput sequencing data in fastq format generated in ChIP-seq experiments and produces lists of genomic loci occupied by the histone modification or bound by the transcription factor under study. For this type of analysis, a parameter file needs to be provided specifying the microalga of interest, execution mode, the fastq files locations or accession numbers, whether an input or mock control sample is included and whether the data has been generated for a histone modification or transcription factor. The results of the pipeline include reports in html and pdf format with information regarding sequence quality analysis, mapping process and identification of the genomic loci occupied by the histone modification or bound by the transcription factor under study. The coordinates of these genomic loci are generated in BED (Browser Extensible Data) format and the genome wide mapping signal is produced in a BigWig (Big Wiggle) file. These formats are supported by ALGAEFUN so that these files can be uploaded directly to proceed with the analysis. To illustrate this pipeline, we re-analysed ChIP-seq data studying the genome wide distribution of the histone modification H3K4me3 in the chlorophyceae microalgae *Chlamydomonas reinhardtii* [24] using the parameter file provided within the MARACAS distribution bundle. The reports produced by MARACAS did not inform of any issue during the processing, leading to the generation of the genomic loci in BED format and the mapping signal in BigWig format.

The graphical interface in ALGAEFUN allows users to input these files to study the genome-wide distribution of specific transcription factors or histone modifications like the H3K4me3 in this case. First, users need to select “Genomic Loci Functional Analysis” from the navigation side panel, Fig. 3(A). Next the microalga of interest has to be chosen from a dropdown menu, *Chlamydomonas reinhardtii* in this example. Users must also set the distance around the transcription start site (TSS) defining gene promoters as well as the gene features that a genomic locus needs to overlap to consider the corresponding gene a target; promoter, 5’,3’-UTR (5’,3’-Untranslated Regions), Exon and Intron, Fig 3(A). The set of genomic loci must be specified by either uploading a BED file or by pasting them in the corresponding text box. Optionally, the genome wide mapping signal can be uploaded in BigWig format, Fig 3(A).

**Figure 3.**
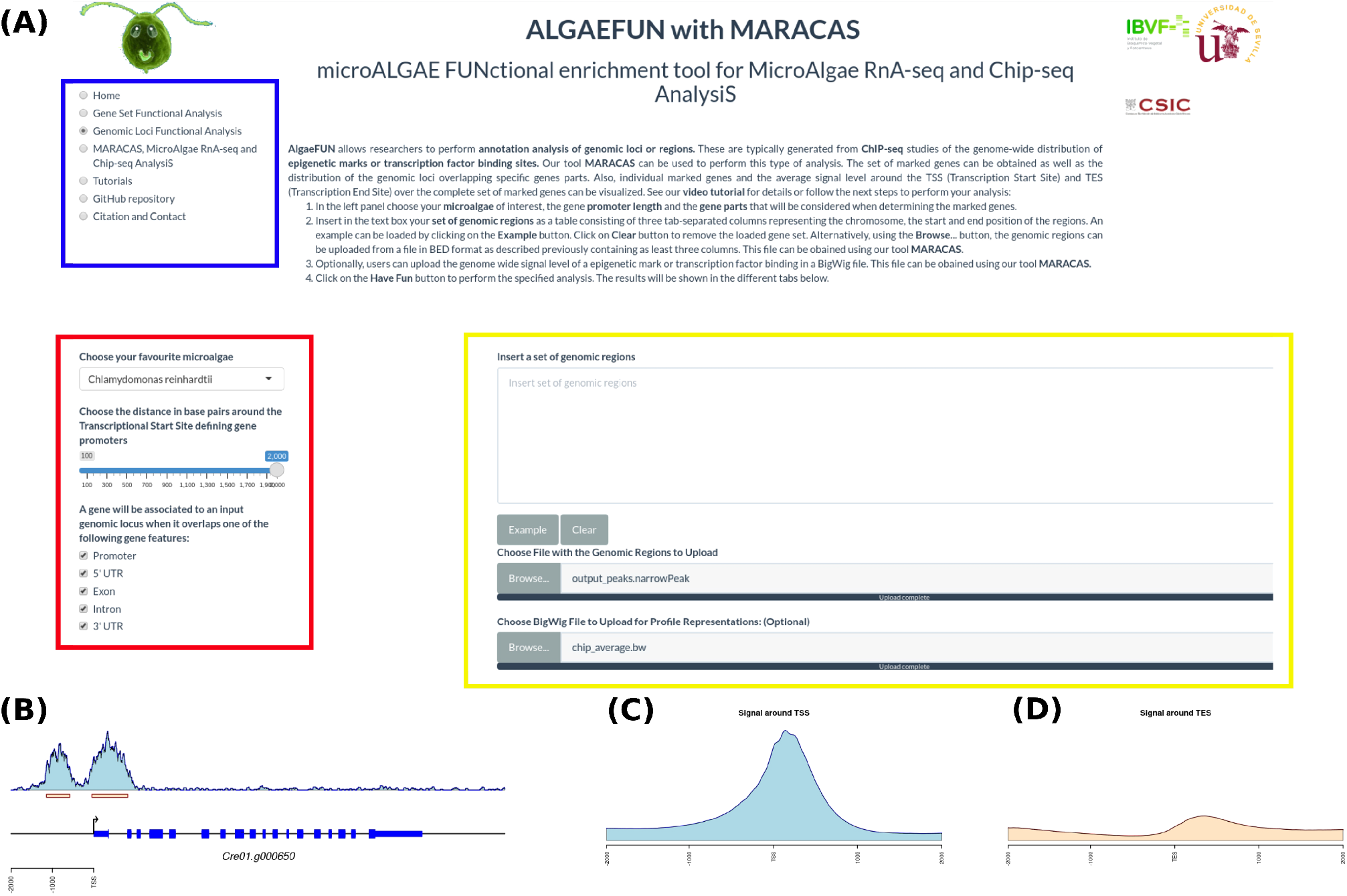
AlgaeFUN Graphical Interface and Results for Genomic Loci Annotation generated from ChIP-seq Analysis. (A) Navigation side panel for selecting the different functionalities (blue box); dropdown menu for microalgae selection and sidebar panel with parameter settings for identifying genes associated with the analysed genomic loci (red box); text box and upload buttons for genomic loci and signal level (yellow box). (B) Visualization of signal distribution over a specific gene and identified peaks in MARACAS. (C) Average signal level around the TSS (Transcription Start Site) for the identified target genes generated as one of the output graphs in AlgaeFUN. (D) Average signal level around the TES (Transcription End Site) for the identified target genes generated as one of the output graphs in AlgaeFUN.

For some microalgae an example consisting of a list of genomic loci can be accessed and inputted in the text box by clicking on the Example button. This would allow users to explore the type of results generated by ALGAEFUN when analysing the outcomes of a ChIP-seq experiment. These examples were generated during the testing of MARACAS using previously published ChIP-seq data sets and have in turn been used in the testing and validation of ALGAEFUN.

In this case study we uploaded to ALGAEFUN the 12,814 genomic loci identified by MARACAS as significantly occupied by H3K4me3 in the *Chlamydomonas* genome under standard growth conditions and the corresponding genome wide mapping signal file in BigWig format. We considered as gene promoter the region two kilobases around the TSS and selected all the gene features to determine the H3K4me3 marked genes. The outputs are presented in the graphical interface in different tabs. A downloadable table with the marked genes and their available annotation is generated. This gene list can in turn be analysed by ALGAEFUN to perform a GO term and/or pathways enrichment analysis. We identified 11,558 H3K4me3 marked genes. Graphs representing the distribution of the genomic loci overlapping different gene features and the distance distribution upstream and downstream from genes TSS are also represented. In agreement with previously published results, we found that 90.75% of the genomic loci occupied by H3K4me3 are located at gene promoters in *Chlamydomonas*. As in this case study, when a BigWig file with the genome wide mapping signal is provided, specific marked genes can be selected to visualize the signal profile over their gene bodies and promoters. A gene example presenting two H3K4me3 peaks on its promoter is depicted to illustrate this functionality in Fig 3 (B). Moreover, DNA motifs recognized by specific transcription factors and regulators in photosynthetic organisms can be identified in the promoter of the selected gene. Finally, a visualization of the average level of signal around Transcriptional Start Site (TSS) and Transcriptional End Site (TES) across all marked genes is generated. For the case of H3K4me3 in *Chlamydomonas* we obtained further evidence showing that this epigenetic mark specifically and exclusively locates at the TSS of marked genes and not at the TES, Fig 3(C,D).

As described above, ALGAEFUN with MARACAS constitutes one of the first steps that has been taken for the development of tools that would enable the microalgae research community to exploit high throughput next generation sequencing data by applying systems biology techniques. For the model microalgae *Chlamydomonas reinhardtii*, researchers can find several online tools to functionally annotate set of genes, such as Algal Functional Annotation Tool [52] and ChlamyNET [53]. Only the online tool AgriGO [54] offers the possibility of analysing a restrictive number of different microalgae species beyond *Chlamydomonas*. However, these tools are only applicable to perform functional annotation of gene sets based exclusively on Gene Ontology (GO) enrichment analysis. The identification of significantly enriched Kyoto Encyclopedia of Genes and Genomes (KEGG) pathways in the inputted sets of genes is not supported. Moreover, none of these tools can be used to process high-throughput sequencing raw data from RNA-seq or ChIP-seq experiments, or functionally annotate genomic loci obtained from a ChIP-seq analysis. In this respect, ALGAEFUN with MARACAS improves and implements several novel functionalities of similar already existing software tools, Table 2.

**Table 2.** Comparison between ALGAEFUN with MARACAS and other functional enrichment analysis tools.

|  | Algal Functional<br>Annotation Tool | AgriGO | ChlamyNET | ALGAEFUN with<br>MARACAS |
| --- | --- | --- | --- | --- |
| Gene Sets as<br>Input | YES | YES | YES | YES |
| Genomic Loci<br>as Input | NO | NO | NO | YES |
| GO<br>enrichment | YES | YES | YES | YES |
| KEGG<br>pathways<br>enrichment | YES | NO | NO | YES |
| Several<br>Microalgae | NO | YES | NO | YES |

## Conclusion

The main contributions of the tool introduced herein, ALGAEFUN with MARACAS, are enumerated below:

1. ALGAEFUN with MARACAS provides a platform specifically designed for microalgae to analyse raw high-throughput sequencing data from RNA-seq and ChIP-seq studies and functionally annotate the resulting genes sets and/or genomic loci. Our goal consists of developing freely available and easy to use tools that would enable the microalgae research community to perform molecular systems biology analysis.
2. In order to interpret the biological relevance of the gene sets and genomic loci obtained from RNA-seq and ChIP-seq data analysis ALGAEFUN with MARACAS provides simple and informative graphical representations of the GO functional and KEGG pathways enrichment results.
3. Most similar annotation tools only support functional enrichment analysis for gene sets and are restricted to the model microalgae *Chlamydomonas*. In contrast, our annotation tool ALGAEFUN also supports results obtained from ChIP-seq analysis and a wide range of different microalgae species.

## Availability and requirements

Project name: AlgaeFUN with MARACAS

Project home page: https://greennetwork.us.es/AlgaeFUN/

Operating system(s): Platform independent

Programming language: Bash, R, shiny

Other requirements: none

License: GPL-3.0 License

## Availability of data and materials.

The code for ALGAEFUN with MARACAS together with the R scripts used in data processing are publicly available at their respective GitHub repositories from the following links: https://github.com/fran-romero-campero/ALGAEFUN and https://github.com/fran-romero-campero/MARACAS. The *Ostreococcus tauri* RNA-seq data set used in the first case study is freely available on the SRA database identified with accession number SRP066656. The *Chlamydomonas reinhardtii* ChIP-seq data set used in the second case study is freely available on the GEO database identified with the accession number GSE59629.

## Abbreviations

AlgaeFUN: (MicroAlgae Functional Annotation Tool)
BED: (Browser Extensible Data)
ChIP-seq: (Chromatin ImmunoPrecipitation sequencing)
EC: (Enzyme Commission)
FDR: (False Discovery Rate)
GO: (Gene Ontology)
HISAT2: (Hierarchical Indexing for Spliced Alignment of Transcripts 2)
HMMER: (Biological sequence analysis using profile Hidden Markov Models)
KAAS: (KEGG Automatic Annotation Server)
KEGG: (Kyoto Encyclopedia of Genes and Genomes)
KOG: (Eukaryotic Orthologous Groups)
LIMMA: (Linear Models for Microarray Analysis)
MACS2: (Model-based Analysis of ChIP-seq 2)
MARACAS: (MicroAlgae RnA-seq and Chip-seq AnalysiS)
RNA-seq: (RNA sequencing)
PANTHER: (Protein Analysis Through Evolutionary Relationships)
PFAM: (Protein Family)
SLURM: (Simple Linux Utility for Resource Management)
TES: (Transcription End Site)
TSS: (Transcription Start Site)
UTR: (Untranslated Region)

## Acknowledgements

This work was supported by the research project MINOTAUR (BIO2017-84066-R) from the Spanish Ministry of Science and Innovation and Intramural Action of the Spanish National Research Council (201420E035).

## Authors’ contributions

ABRL, FJRC, CA and PdlR co-developed all the components of MARACAS, ALGAEFUN and the different annotation R packages. FJRC and MGG selected the microalgae species to be included in ALGAEFUN with MARACAS, designed the application interface and wrote the manuscript. All authors read and approved the manuscript.

## Declarations

Ethics approval and consent to participate. Not applicable

Consent for publication. Not applicable

